# Cyclic peptides space: The methodology of sequence selection to cover the comprehensive physical properties

**DOI:** 10.64898/2026.03.10.710724

**Authors:** Ryo Tsuchihashi, Misaki Kinoshita

## Abstract

Cyclic peptides have emerged as a pivotal modality for next-generation therapeutics, due to their superior biocompatibility, high selectivity, and structural stability. While AI-driven peptide design has advanced rapidly, conventional optimization algorithms are often constrained by initialization biases, which impede the efficient exploration of the vast chemical space. Here, we propose a novel methodology that integrates the protein language model ESM-2 with cyclic permutation averaging of embeddings to resolve this bottleneck. This approach establishes a comprehensive "peptide space", a high-dimensional vector representation that encapsulates the physicochemical and structural attributes of cyclic peptides. Our analysis reveals that random sequence selection results in a heterogeneous distribution within this space, potentially underrepresenting specific functional regions. Conversely, navigating this defined peptide space enables the selection of libraries that uniformly span diverse molecular properties. In a proof-of-concept study designing binders for β2-microglobulin (β2m), we demonstrate that initial sequences uniformly sampled from our peptide space yield superior candidates more efficiently than those derived from random selection. Furthermore, this framework facilitates the quantitative assessment of mutational perturbations on global peptide properties, supporting rational decision-making for both broad exploration and local optimization. This "peptide space" concept provides a foundational framework for defining appropriate search boundaries and enhancing computational efficiency in AI-mediated drug discovery.

**Graphic Abstract:** 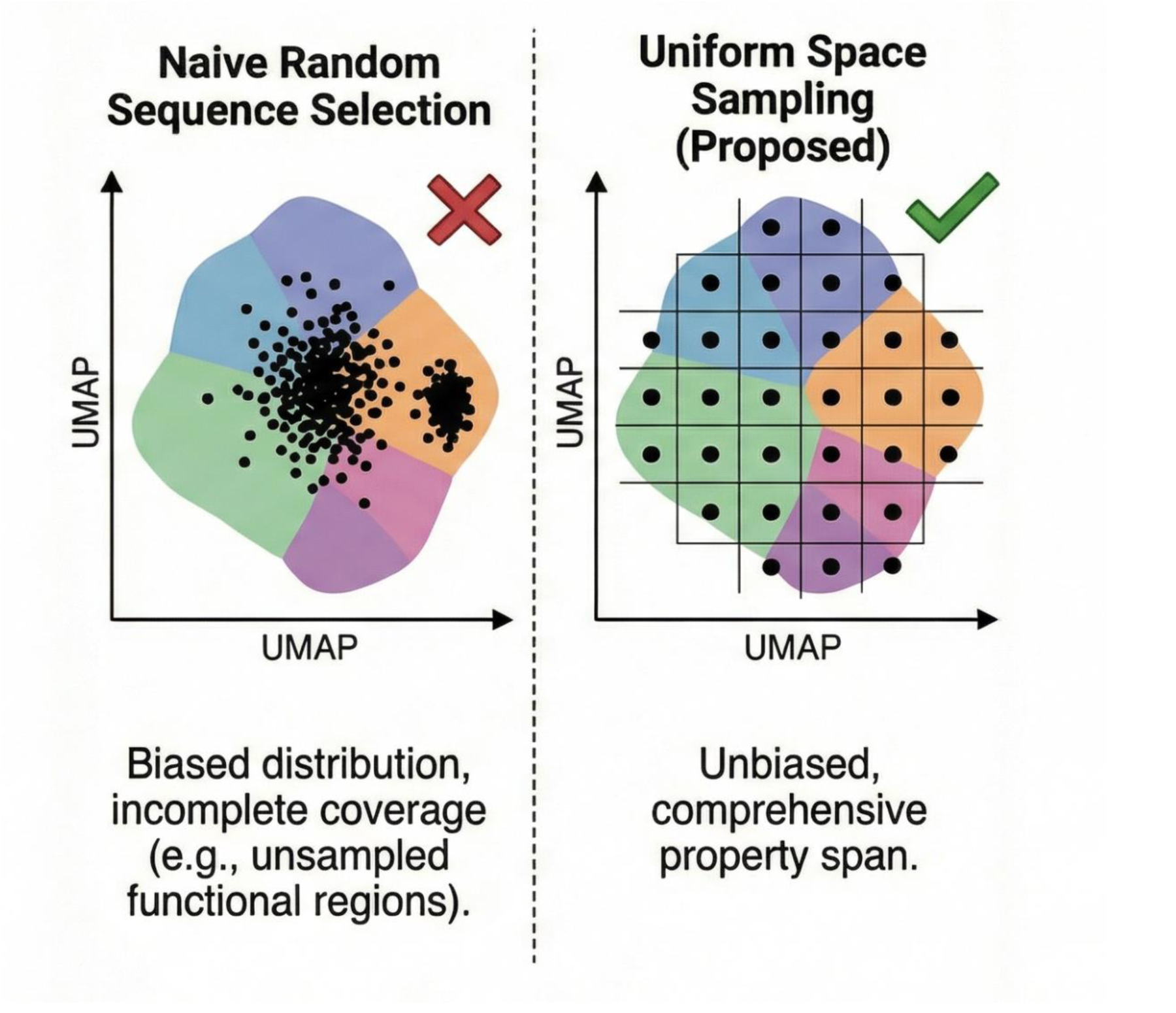

## Introduction

In the landscape of pharmaceutical development, peptide therapeutics have emerged as a pivotal modality, bridging the gap between conventional small molecules and antibody biologics [1, 2]. Peptides offer a unique advantage by accessing complex chemical spaces and adopting intricate conformations, thereby enabling high-affinity interactions with a broad spectrum of disease-associated proteins which are often considered "undruggable" by small molecules. Furthermore, they can be engineered for specific properties such as enhanced tissue penetration. Compared to antibodies, peptides possess superior biocompatibility due to their construction from fundamental amino acids and offer the distinct benefit of cost-effective synthetic manufacturability [3, 4]. Consequently, the market for peptide therapeutics is expanding rapidly, with successive approvals in diverse therapeutic areas including diabetes, oncology, cardiovascular diseases, and rare disorders [5]. Among these, cyclic peptides represent a privileged scaffold. Beyond the general advantages of ease of synthesis and high target selectivity, macrocyclization confers critical physicochemical benefits: resistance to peptidases (proteolytic stability), reduced conformational entropy leading to higher binding affinity, and improved membrane permeability [3]. While recent advancements in AI-driven design have significantly enhanced the accuracy and productivity of cyclic peptide engineering, the comprehensive exploration of the vast combinatorial search space remains computationally prohibitive [6]. To mitigate this, existing computational frameworks employ heuristic optimization algorithms to accelerate the search process. For instance, recent systems such as HighPlay [7] and EvoBind2 [8] utilize evolutionary algorithms initiated from arbitrarily determined sequences. However, because the initial seed sequences profoundly influence the trajectory and quality of the final solution in these stochastic methods, defining an appropriate "search space" for initialization is crucial. Yet, systematic approaches to define such spaces have been lacking. Although some approaches, like RFpeptide, attempt to address this by utilizing a predefined initial space of 1,200 sequences rather than a single seed [9], this space is constructed with a primary focus on secondary structure characteristics, largely overlooking the diversity of chemical properties.

In this study, we report the construction of a comprehensive design space that facilitates the rational determination of initial sequences, incorporating a broader range of physicochemical and structural attributes. We generated a high-dimensional vector space by transforming random amino acid sequences using the protein language model ESM-2 [10]. We evaluated this space for its uniformity in physicochemical properties and its utility in binder design. Our results demonstrate that this peptide space enables the unbiased selection of properties and serves as a powerful tool for optimizing both initial sequence selection and evolutionary directionality. This framework provides a novel methodology for the efficient and appropriate selection of cyclic peptide sequences through the explicit definition and understanding of the search space.

## Methods

### Protein Language Model and Embedding Generation

To extract numerical feature vectors (embeddings) from peptide sequences, we employed the pre-trained protein language model ESM-2 (esm.pretrained.esm2_t6_8M_UR50D) [10]. Specifically, we utilized residue-level representation vectors derived from the intermediate layer (layer 6) [11]. To ensure deterministic and reproducible embeddings, the model was operated in evaluation mode, thereby disabling dropout layers.

### Representation Vectors for Cyclic Peptides

Cyclic peptides lack a defined N- or C-terminus. To incorporate this topological characteristic into the vector representation, we introduced a "Cyclic Permutation Averaging" strategy. For a peptide sequence *S* of length *L*, we first generated all *L* possible cyclic permutations (denoted as sequence set *S*_*i*_, where *i* = 0, …, *L* − 1, and *S*_0_ is the original sequence), effectively shifting the sequence one residue at a time:

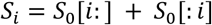

Next, for all *L* sequences, we computed the representation vector *R*_*i*_ for the entire sequence using the ESM-2 model described above. We calculated the arithmetic mean of these *L* sequence representation vectors to obtain the topology-invariant embedding vector, *R*_*cyclic*_, for the original cyclic peptide *S*:

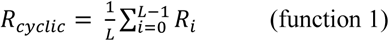

where *R*_*i*_ indicated the vector for the sequence *S*_*i*_. As a control, we also defined a conventional linear peptide representation derived from a single sequence, without applying cyclic permutation averaging.

### Construction of Peptide Space and Dimensionality Reduction

In this study, we focused on peptides with a length of 14 amino acids. We constructed a large-scale random peptide library consisting of approximately one million (*N*∼300,000) sequences, generated by randomly sampling from the standard 20 amino acids. High-dimensional embedding vectors were calculated for all sequences using the Cyclic Permutation Averaging method defined above. To visualize and analyze the global structure of this high-dimensional embedding landscape, we applied Uniform Manifold Approximation and Projection (UMAP), a non-linear dimensionality reduction technique [12]. UMAP was used to project the high-dimensional vectors onto a two-dimensional plane. In this manuscript, we refer to this 2D projection as the "Peptide Space."

### Characterization of the Peptide Space

To characterize the landscape of the constructed chemical space, we performed the following three analyses. First, we quantified the distribution density of peptide sequences within the peptide space using Kernel Density Estimation (KDE) to visualize potential sampling biases. For bandwidth selection, we employed Scott’s Rule, which automatically optimizes the parameter based on sample size and dimensionality to achieve a balance between preventing over-smoothing and avoiding noise over-fitting. Second, The abundance of specific amino acid residues was mapped onto the peptide space. This allowed us to evaluate the correlation between intrinsic chemical properties and their relative positioning within the projected manifold. Finally, to assess the local and global organization of the space, we generated two derivative sets from reference sequences: "shuffled sequences" (preserving composition but altering order) and "variant libraries" (comprising comprehensive residue substitutions). We then analyzed their spatial distribution and calculated the cosine similarity within the cyclic peptide space to define the relationship between sequence homology and spatial proximity.

### Empirical Study: Optimization and Sampling Strategy for β2m binders

To demonstrate the utility of our approach, we analyzed datasets obtained from computational optimization simulations of binder peptides against β2m. To evaluate the impact of initial seed selection on discovery efficiency, we compared two distinct sequence set. One is the Systematic sampling set, in which the peptide space was partitioned into a uniform grid. We identified 92 specific grids that met a minimum sequence density threshold. Representative sequences were then uniformly extracted from each of these grids to ensure maximal coverage of the manifold. Another is the Random sampling set. In this set, sequences were extracted from the library via stochastic selection, without regard to their spatial distribution or chemical diversity. To ensure a rigorous comparison, the total number of sequences for both sequence set was fixed at 920, corresponding to 10 sequences per grid. Using these seed sequences, we conducted binding discovery and structure prediction simulations employing EvoBind2. The primary metric was the Loss value, a composite score reflecting predicted binding free energy and structural stability (Equation 2), where lower values indicate higher predicted affinity. Simulations were organized into 10 independent sets, each comprising 92 sequences (corresponding to the number of grids). The minimum Loss value from each set was extracted to generate box plots for comparative statistical analysis.

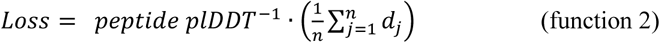

### Correlation between Peptide Space and Structure/Properties

We evaluated the correlation between coordinates in the peptide space and the structural features and physicochemical properties of the peptides. Structural predictions were performed using AfCycDesign [15] for 3,000 sequences randomly selected within the peptide space. From the predicted structures, we calculated topological features—such as radius and inter-atomic distances—secondary structure content, and known physicochemical properties, including hydrophobicity and charge.

## Results and Discussion

### Construction of a cyclic peptide space for unbiased property selection

Establishing an appropriate initialization strategy for cyclic peptide design requires the construction of a search space that is sufficiently discrete and representative of diverse physicochemical and structural properties. We utilized the protein language model ESM-2 [10] to transform randomly generated sequences into high-dimensional vectors that encapsulate their inherent characteristics. Given that ESM-2 is trained on linear sequences, we addressed the cyclic nature of our targets by generating cyclic permutations of the amino acid sequence (shifting the starting position sequentially), computing embeddings for each permutation, and averaging them to derive a definitive "cyclic peptide vector" (Fig. 1A). We constructed vector spaces for datasets comprising 1,000, 10,000, and 300,000 random 14-residue sequences. Dimensionality reduction via UMAP [12] revealed that spaces exceeding 10,000 sequences exhibit a characteristic distribution partitioned into three distinct segments. Notably, random sampling resulted in significant heterogeneity in occurrence frequency within this peptide space (Fig. 1A, right panels). While similar segmentation is observed in linear sequences (Supporting Figure 1), the cyclic vectors derived from the 300,000-sequence dataset eliminated several minor clusters observed in linear counterparts, suggesting that our cyclic permutation averaging effectively smooths out subtle variances caused by linear edge effects.

**Fig. 1.**
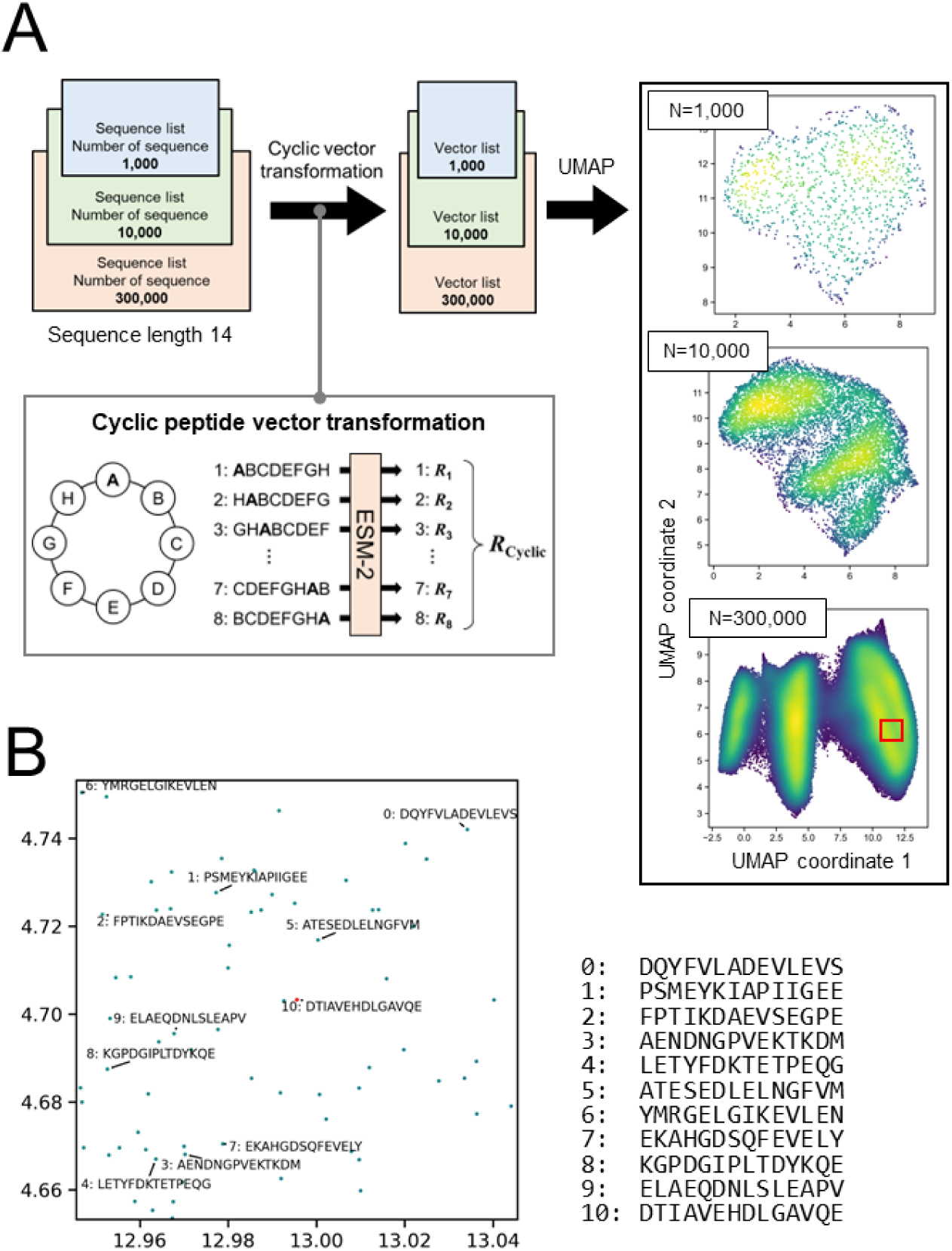
Construction and characterization of a latent space for cyclic peptides. **(A)** Schematic workflow for generating cyclic peptide embeddings. Randomly generated peptide sequences (length 14) with varying dataset sizes (*N* = 1,000, 10,000, and 300,000) were transformed into high-dimensional vectors using the ESM-2. To accommodate the circular topology, the final representation (*R*_*Cyclic*_) was computed by aggregating embeddings derived from all cyclic permutations of the linear sequence (inset). Dimensionality reduction via UMAP (right panels) reveals that a characteristic tripartite distribution emerges as the sample size exceeds 10,000. **(B)** Local analysis of sequence-space relationships. A magnified view of the high-density region (indicated by the red square in **A**) shows the projection of specific peptide sequences with sequence list (right) corresponding to the scatter points. Since the peptide space is defined by the UMAP-based coordinate system, the x and y axis labels are omitted.

To validate the robustness of the proposed cyclic permutation averaging method, we evaluated the variance and cosine similarity of four vector sets derived from a single seed sequence (DTIAVEHDLGAVQE): (i) linear vectors from cyclic permutations; (ii) cyclic vectors from cyclic permutations; (iii) linear vectors from shuffled sequences; and (iv) cyclic vectors from shuffled sequences (Supporting Figure 2). For the linear vectors without averaging (i), slight fluctuations were observed depending on the starting amino acid position, resulting in a cosine similarity of 0.997764 (Supporting Figure 2B, C). This bias is attributed to edge effects (N/C-terminal recognition) inherent in linear models. In contrast, the vectors obtained through the cyclic permutation averaging method (ii), converged to nearly identical coordinates in the peptide space regardless of the input permutation, achieving a perfect cosine similarity of 1.0000. Although these vectors occasionally appeared slightly separated into two points on the UMAP plot due to computational noise or approximation, they were confirmed to be effectively identical. Conversely, the shuffled sequences (iii, iv)—which maintain the same amino acid composition but randomized order—showed significant dispersion in the space for both linear and cyclic formats (Supporting Figure 2B, C). These results demonstrate that the vectors generated by our method reflect structural and chemical properties based on sequence order rather than simple composition, thereby appropriately representing the topology of cyclic peptides.

**Fig. 2.**
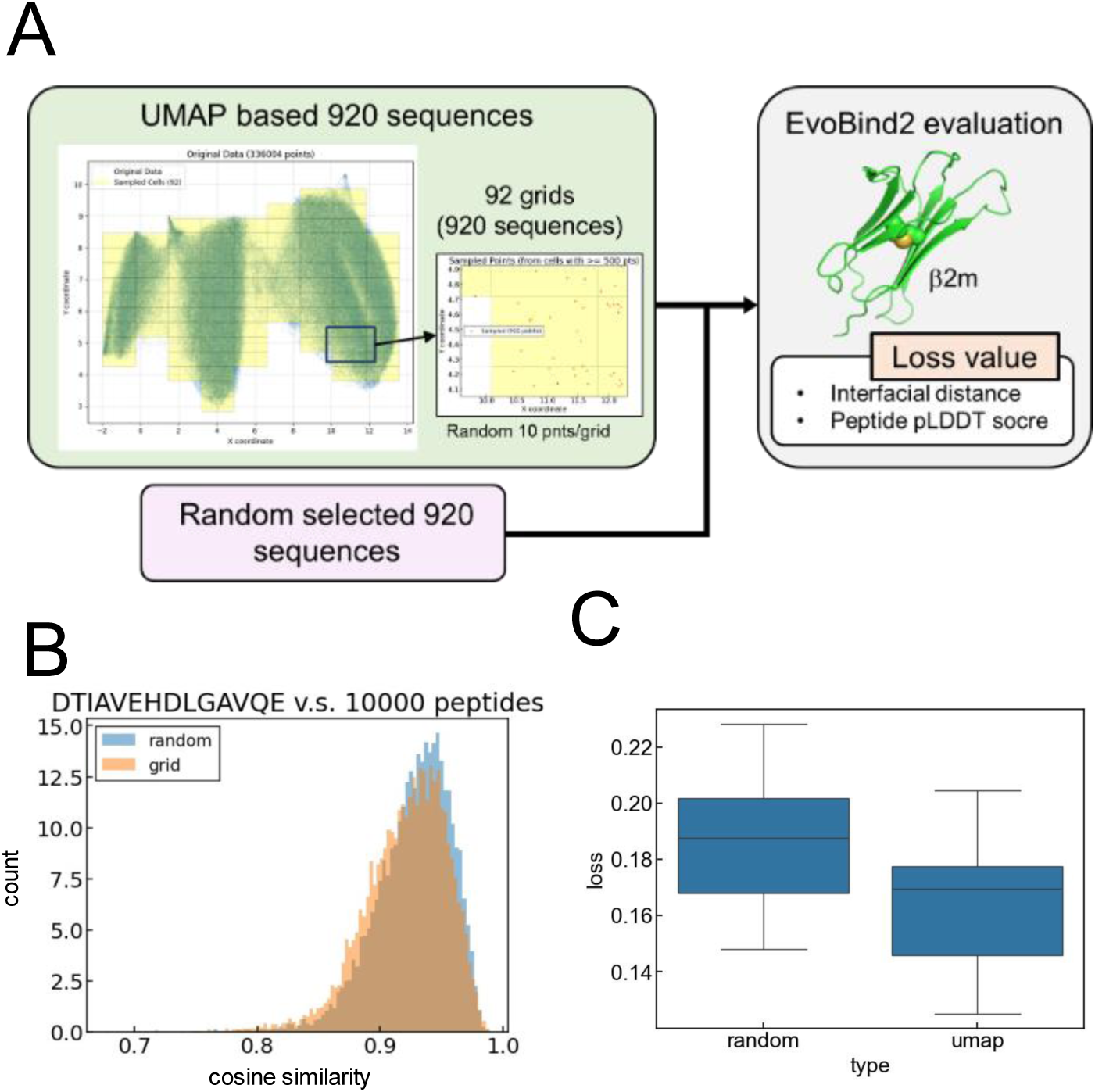
Enhanced efficiency of cyclic peptide design via latent space-guided initialization. **(A)** Workflow for evaluating the utility of the constructed peptide space in targeted design. The target protein, β2m, was used to test the generation of cyclic peptide binders via EvoBind2 [8]. To mitigate sampling bias, the peptide space was segmented into 92 discrete grids, and 10 sequences were randomly drawn from each to create a structurally diverse "UMAP-based" dataset (N=920). This was compared against a control "Random" dataset (N=920) generated stochastically. Loss values, calculated based on interfacial distance and peptide pLDDT scores (inset), served as the evaluation metric. **(B)** Assessment of sequence diversity. The histogram displays the distribution of cosine similarities for the UMAP-based (orange) versus random (blue) datasets against a reference sequence. The grid-based selection exhibits a broader distribution with a higher frequency of lower similarity scores, indicating that spatial segmentation effectively reduces redundancy and captures a wider range of physicochemical properties. **(C)** Comparison of design performance. Box plots representing the distribution of loss values after EvoBind2 optimization.

Mapping randomly picked sequences onto this space revealed no correlation between spatial distance and sequence similarity, indicating that the space captures properties independent of simple sequence homology (Fig. 1B). Furthermore, to assess the "randomness" relative to a specific query, we calculated cosine similarities between the seed sequence (DTIAVEHDLGAVQE) and 10,000 random sequences (Fig. 2B). The majority of random sequences clustered within a high similarity range (0.85–1.00), demonstrating that naive random sampling frequently results in vectors with redundant properties. Color-coding the distribution by amino acid count revealed that the three major segments in the peptide space are largely defined by cysteine content (Supporting Figure 3A). Other residues also drive spatial bias; for instance, peptides constructed with more than two methionine localize between the central and right segments, while those with constructed with more than two tryptophan show distinct biases within segments. This underscores that simple amino acid composition significantly skews the distribution.

**Fig. 3.**
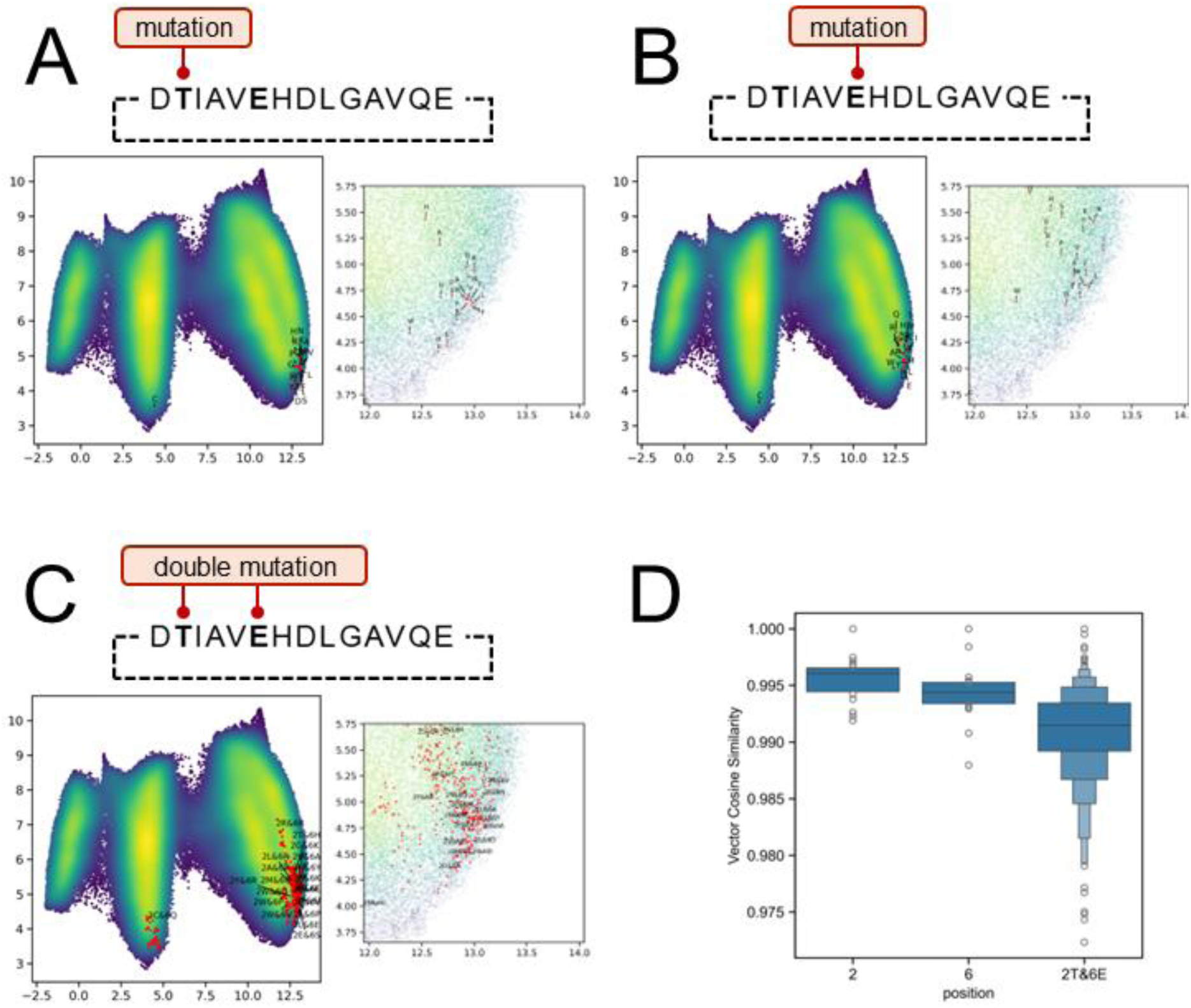
Visualization and quantification of mutational effects in the latent space. (A–C) Projection of mutant sequences onto the peptide space. The reference sequence was subjected to exhaustive point mutations at position 2 (A), position 6 (B), or simultaneous mutations at both positions (C). In each panel, the left plot displays the global distribution of the resulting mutant vectors within the constructed UMAP landscape, while the right plot provides a magnified view of the local region surrounding the parental sequence to visualize the relative positioning of individual variants. (D) Distribution of vector similarities. Box plots show the cosine similarity of single ("2" and "6") and double ("2T&6E") mutant vectors relative to the reference sequence. Since the peptide space is defined by the UMAP-based coordinate system, the x and y axis labels are omitted.

We further analyzed structural attributes by generating 3D structures for 1,000 randomly selected sequences using AfCycDesign [15], an AlphaFold-based tool for cyclic peptides. We mapped various structural or physical features onto the peptide space (Supporting Figure 3B). Rather than a uniform distribution, these parameters exhibited a mosaic-like, patchy organization. Physicochemical properties showed clear biases, consistent with the aforementioned compositional skew. Similarly, secondary structure and conformational metrics displayed non-uniform, localized patterns. Collectively, these findings suggest that random sequence selection fails to guarantee physicochemical or structural stochasticity, potentially making it an unsuitable strategy for initialization. Our proposed "peptide space" offers a solution to overcome these biases by enabling rational, uniform sampling.

### Use case: Enhanced binder design via peptide space navigation

To demonstrate the utility of this peptide space, we applied it to the design of cyclic peptide binders for β2m using EvoBind2 [8]. β2m is a component of MHC class I molecules, essential for presenting self-antigens to cytotoxic T cells [16]. With its seven-stranded `β`-sandwich fold and single disulfide bond within a compact 99-residue structure, β2m represents an ideal target for evaluating complex prediction. To ensure uniform property sampling, we segmented the peptide space into 92 grids and randomly selected 10 sequences from each, creating a "UMAP-based sequence set" (920 sequences). A control "Random sequence set" of equal size was generated stochastically (Fig. 2A). The optimization process involved 10 iterations, each using a subset of 92 sequences (one from each grid for the UMAP-based set, and a random subset for the control).

The grid-based selection demonstrated a broader distribution of cosine similarities compared to random selection, specifically increasing the probability of sampling sequences with lower similarity (i.e., unique properties) (Fig. 2B). We then performed EvoBind2 design targeting β2m for each subset and calculated the Loss value using the equation 2 described in the methods section. The UMAP-based sequence set yielded consistently lower mean and minimum Loss values compared to the Random sequence set, indicating a higher success rate in identifying computationally favorable candidates (Fig. 2C). Furthermore, mapping the loss values from the UMAP-based set in the peptide space revealed that the sequences exhibiting the lowest loss values are located at the segment boundaries (Supplementary Figure 4), because this region is often missed by random sampling due to its statical rarity. These results demonstrate that navigating this peptide space ensures true physicochemical coverage, which enabled the efficient identification of high-potential candidates that would otherwise be overlooked.

### Quantifying mutational effects on peptide properties

Finally, we utilized this space to quantitatively evaluate how specific amino acid substitutions perturb global peptide properties. Using the sequence DTIAVEHDLGAVQE, we substituted the residues at positions 2 and 6 with all 20 amino acids and analyzed the resulting shifts in the peptide space and cosine distances. Single substitutions at either position caused minimal spatial displacement, with the notable exception of cysteine (Fig. 3A, B). Cysteine introduction triggered a segment shift in peptide space, reflecting the dominant influence of disulfide potential (Supporting Figure 3A). While non-cysteine mutations showed no consistent directional trend in peptide space (Fig. 3A, B, right panels), hierarchical clustering based on cosine similarity revealed clear physicochemical logic (Supporting Figure 5). This discrepancy arises because UMAP dimensionality reduction preserves local topology but may distort global directional relationships. Clustering analysis confirmed that substitutions with chemically similar residues—such as Asp/Glu, Ser/Thr, Tyr/Phe/Ile, and Ala/Val—formed tight clusters with high similarity. This is intuitive given their side-chain properties. Interestingly, basic residues (Lys, Arg, His) did not form a unified cluster, indicating that in the context of cyclic peptides, these residues impart distinct vector characteristics, likely due to differences in functional group geometry and hydrophobic surface area. Double mutations (positions 2 and 6) resulted in a synergistic expansion of distribution in peptide space and high variance in cosine similarity (Fig.3C, D). This highlights two key capabilities: (1) the ability to switch segments by altering cysteine content, and (2) the ability to fine-tune similarity within a segment by combining specific mutations. In conclusion, this peptide space provides a powerful framework for rational design, enabling users to select mutations that match their specific exploration goals—whether it be a broad "jump" to a new physicochemical regime or a local "optimization" preserving the parent character.

## Conclusion

In this study, we proposed the novel concept of "Peptide Space." This framework employs a strategy where linear sequence embeddings generated by ESM-2 are averaged over a set of cyclic permutations, explicitly designed to mitigate the bias of terminal effects inherent in language models trained on linear proteins. Our analysis demonstrated that naive random selection of sequences results in a non-uniform distribution of physicochemical and structural properties within the ESM-2 space. This finding underscores a critical discrepancy in conventional machine learning-based design: while the objective of initialization is to cover a "random" spread of physicochemical properties, relying on "sequence" randomness leads to the systematic undersampling of low-frequency, yet potentially functional, regions of the chemical space. Our methodology resolves this bottleneck, enabling a truly unbiased initialization that encompasses a broad spectrum of molecular attributes.

In our proof-of-concept study designing binders for β2m using EvoBind2, we compared the binding potential, quantified by Loss values, of candidates derived from our peptide space against those from a random sequence set. The results indicated that navigating the peptide space allows for the identification of superior starting points. This observation likely extends beyond β2m to other therapeutic targets; reliance on stochastic sequence initialization carries a high probability of overlooking sequences with favorable theoretical binding probability (low Loss values), thereby compromising the quality of the final design and inflating computational costs. Conversely, utilizing our peptide space facilitates the selection of appropriate initial seeds, helping to circumvent local minima, enhance the quality of the final design, and reduce the overall computational burden.

From the perspective of computational efficiency, this framework is particularly advantageous for modulating evolutionary directionality. By utilizing changes in the direction and distance of peptide sequence vectors within reduced-dimension coordinates, it becomes possible to rationally select next-generation mutations tailored to specific local optimization goals. For instance, in purpose of excluding mutations that result in negligible vector shifts involving chemically minor changes, efficiency can be significantly improved by pruning the search space rather than relying on blind stochastic mutation. Although integrating ESM-2 calculations adds a step to the peptide design workflow, the inference time is minimal (<1 ms per sequence [10]) and is far outweighed by the efficiency gains achieved through the strategic reduction of the search space.

A fundamental insight of this work is the critical importance of comprehending the nature of the search space itself. As the application of machine learning expands from cyclic peptides to broader domains of drug discovery and materials science—including the design of protein binders and inhibitors [17]—the associated computational costs are becoming a non-negligible barrier. Our approach, while conceptually simple, demonstrates that explicitly defining and understanding the search space can lead to substantial cost reductions and more rapid, robust design cycles. When the concept of "Chemical Space" was first introduced, its primary utility was the comprehensive coverage of features [18, 17]. However, in the current era of generative AI, we argue that this concept has evolved to play a dual role: ensuring diversity and optimizing computational tractability. As new machine learning architectures for peptides and proteins continue to emerge, methods that provide a structured understanding of the exploration landscape—such as the one proposed here—will play an indirect but indispensable role in maximizing the efficiency of our limited computational resources.

## Acknowledgements

-

## Statements and Declarations

### Ethics Approval

This article does not contain any studies with human participants or animals performed by any of the authors.

### Data Availability

All data generated or analyzed during this study are included in this published article and its supplementary information files.

## Supplementary information

### Supporting figures

**Supporting Figure 1.**
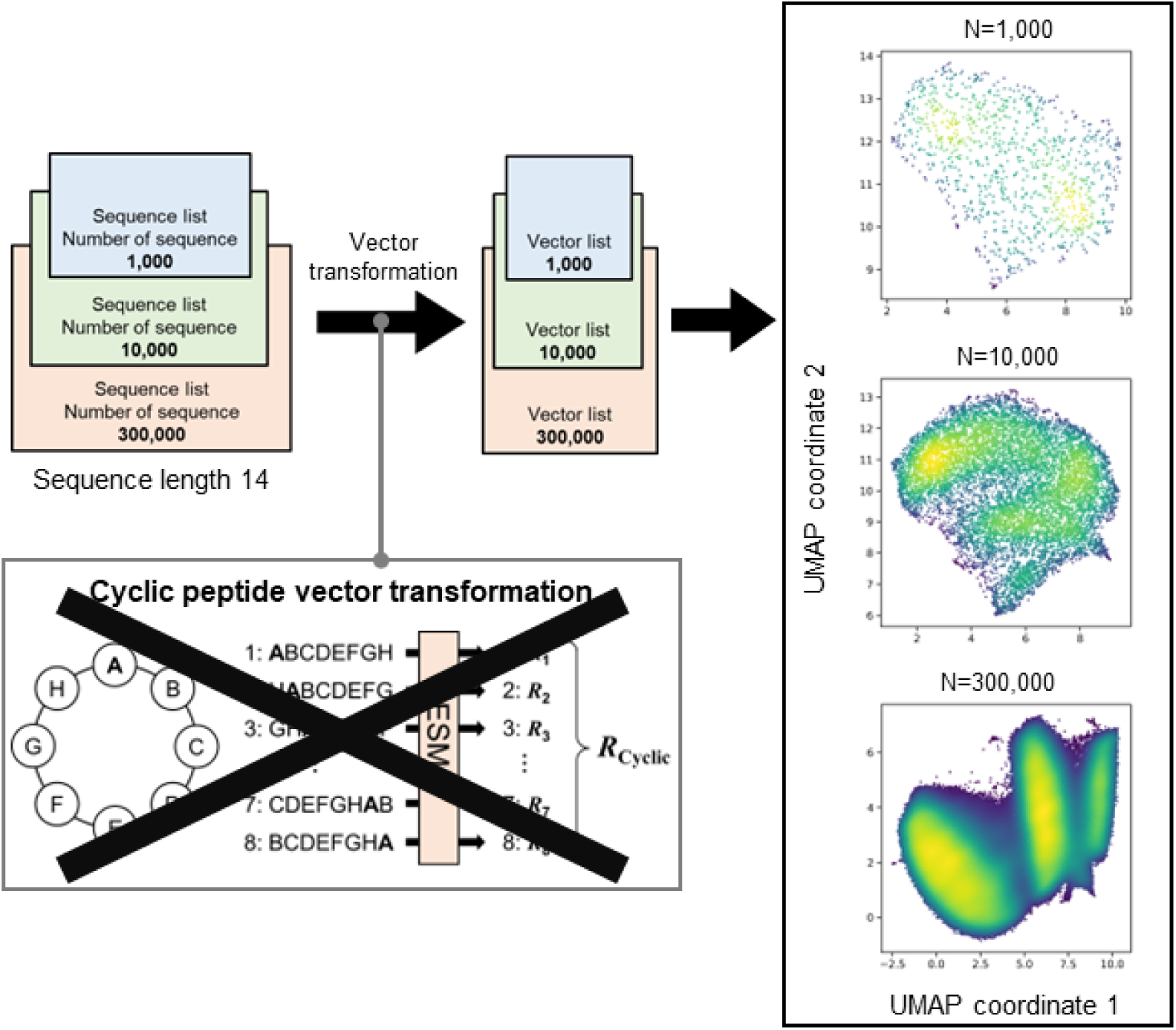
UMAP projection of linear peptide embeddings. Randomly generated 14-residue sequences were encoded using the ESM-2 protein language model. Unlike the cyclic vectorization strategy (Fig. 1A), the cyclic permutation averaging step was omitted (indicated by the crossed-out schematic), and sequences were processed as linear inputs. The resulting high-dimensional vectors were projected into a two-dimensional space using Uniform Manifold Approximation and Projection (UMAP). Density plots are displayed for datasets containing 1,000(upper), 10,000 (middle) and 300,000 (bottom) sequences.

**Supporting Figure 2.**
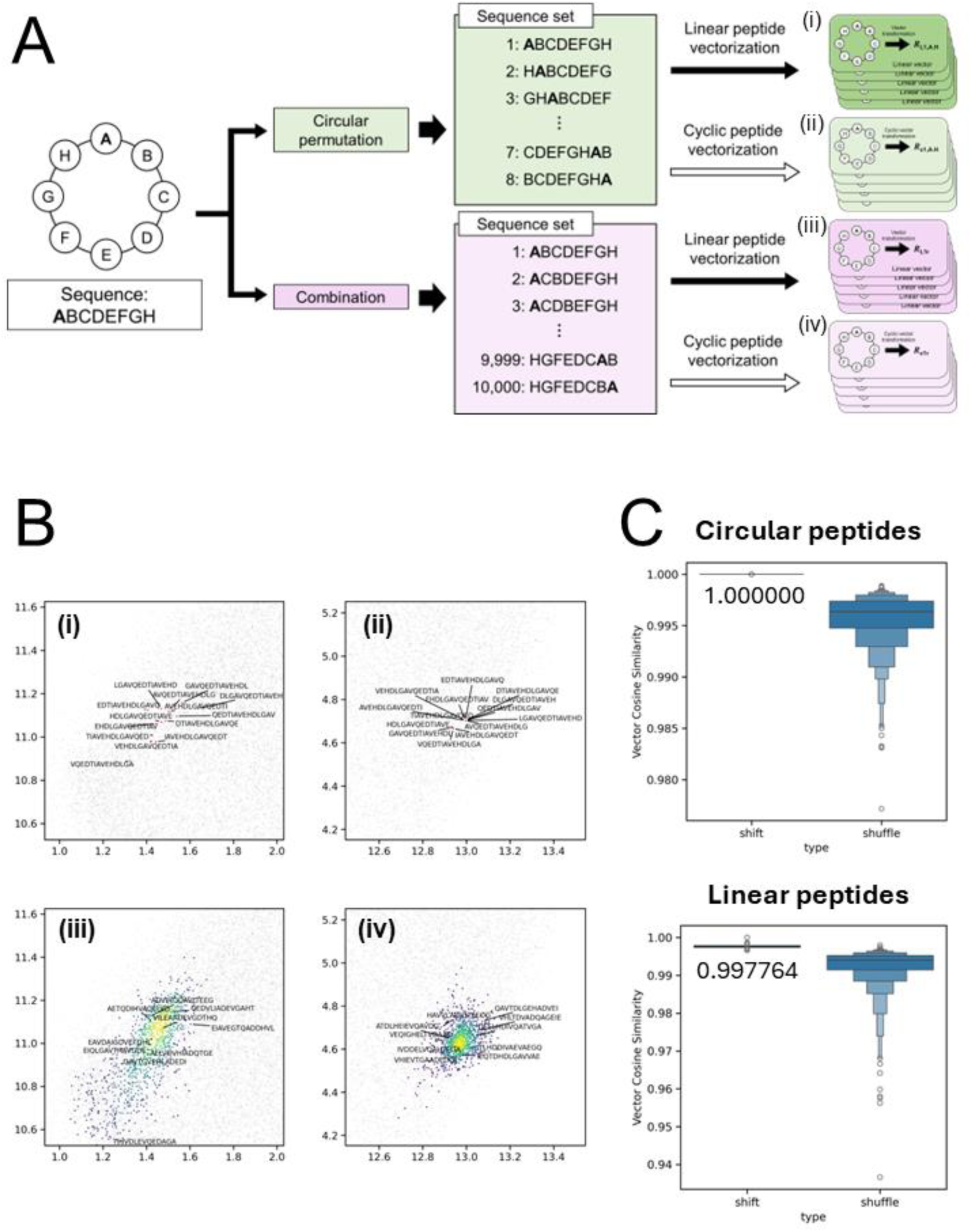
Validation of permutation invariance and sequence specificity in cyclic peptide embedding. **(A)** Experimental design for assessing vector robustness. A seed sequence (DTIAVEHDLGAVQE) was used to generate four distinct datasets to disentangle topological effects from compositional variance: linear and cyclic vectorization applied to (i, ii) 14 cyclic permutations (sequence shifts) and (iii, iv) 10,000 randomized permutations (shuffled combinations). **(B)** Visualization of vector convergence. UMAP projections of the four datasets. Notably, cyclic vectors derived from cyclic permutations (panel **ii**) converge to a single point, visually demonstrating that the embedding is independent of the starting residue. In contrast, linear vectors (panel **i**) exhibit slight spatial divergence due to N- and C-terminal edge effects. Both shuffled datasets (panels **iii** and **iv**) show broad spatial distribution, confirming that the embedding captures sequence order rather than mere amino acid composition. Since the peptide space is defined by the UMAP-based coordinate system, the x and y axis labels are omitted. **(C)** Quantification of embedding stability. Box plots of vector cosine similarities relative to the seed sequence. The cyclic vectorization method achieves perfect stability (similarity = 1.0000) across all cyclic permutations, effectively neutralizing the floating-point divergence observed in linear processing (similarity = 0.9991). The marked variance in shuffled sequences further validates that the vector representation is strictly determined by the specific sequential arrangement of residues.

**Supporting Figure 3.**
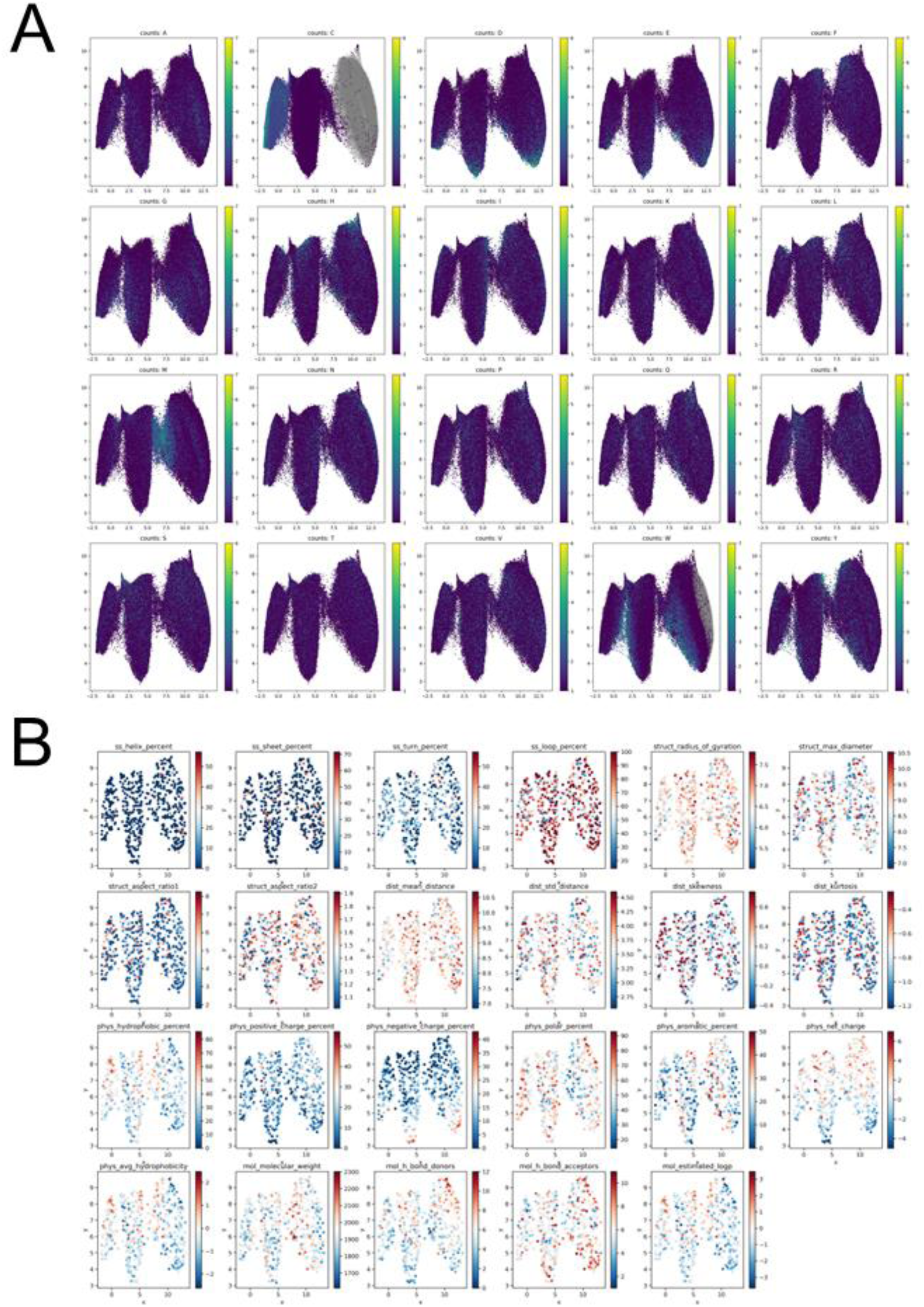
Deconvolution of compositional and structural determinants in the cyclic peptide latent space. **(A)** Mapping of amino acid abundance distributions. The UMAP landscape is color-coded by the count of individual amino acid types within the constituent sequences. Cysteine content (panel ’counts: C’) acts as the primary driver of the global topology, dictating the segmentation into three distinct clusters. Other residues, such as methionine (M) and tryptophan (W), also exhibit specific localized biases, highlighting that simple compositional variations significantly skew the spatial distribution. **(B)** Landscape of structural and physicochemical properties. To correlate the latent space with physical attributes, 3D structures for 3,000 randomly sampled sequences were predicted using AfCycDesign [15]. Various metrics—including secondary structure content (e.g., helix/sheet percentage), geometric parameters (e.g., radius of gyration), and physicochemical nature (e.g., hydrophobicity, charge)—were projected onto the manifold. The resulting "mosaic-like" patterns demonstrate that these properties are not uniformly distributed but are instead organized into discrete regimes. This heterogeneity underscores that stochastic sequence generation fails to guarantee uniform sampling of the structural and physicochemical property space. Since the peptide space is defined by the UMAP-based coordinate system, the x and y axis labels are omitted.

**Supporting Figure 4.**
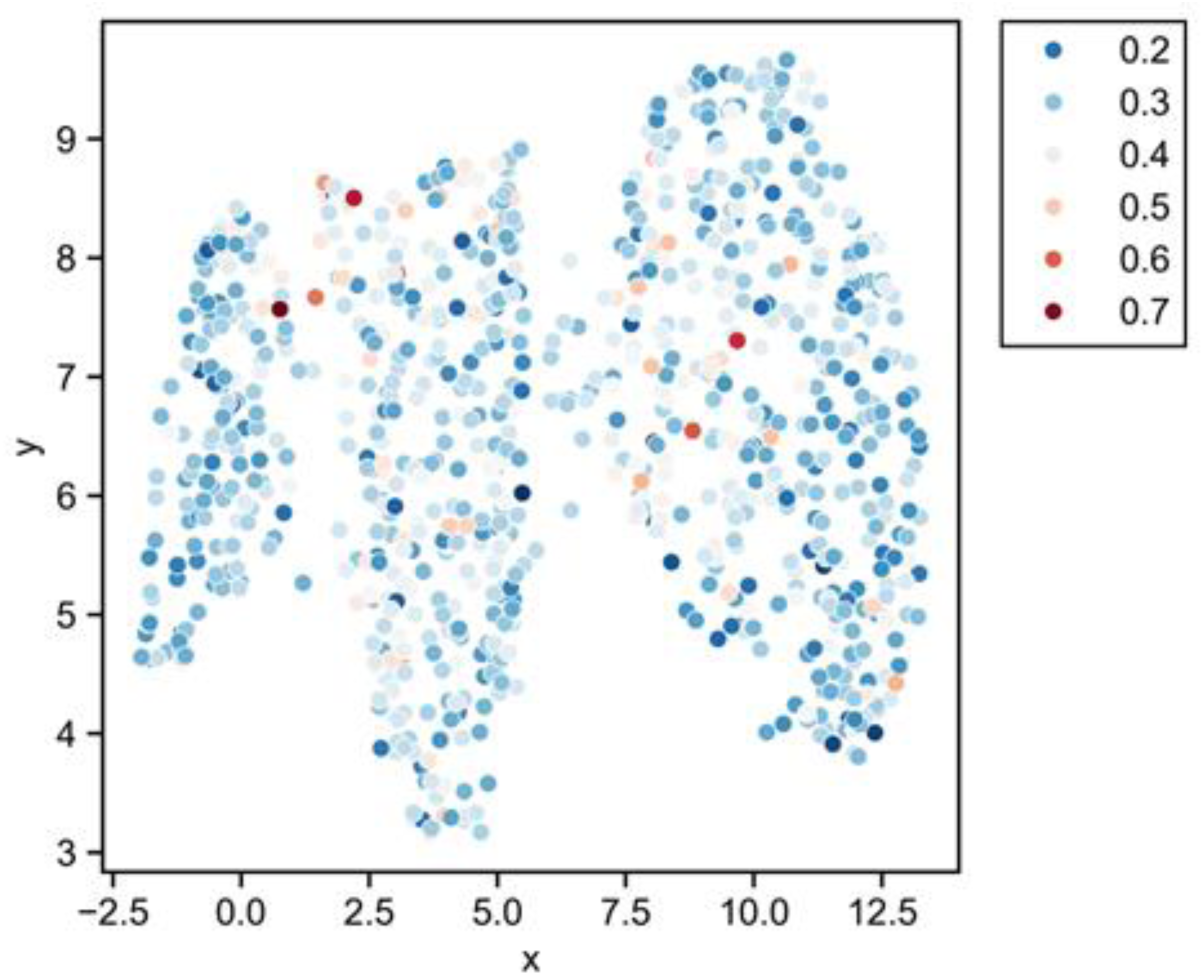
Spatial localization of high-potential binder candidates. Visualization of design performance landscapes. The Loss values derived from EvoBind2 optimization targeting β2m are mapped onto the corresponding coordinates of the starting sequences in the peptide space. The heatmap color scale represents the raw Loss value, where specific points (e.g., darker hues) indicate sequences with superior predicted binding properties (lower loss). Since the peptide space is defined by the UMAP-based coordinate system, the x and y axis labels are omitted.

**Supporting Figure 5.**
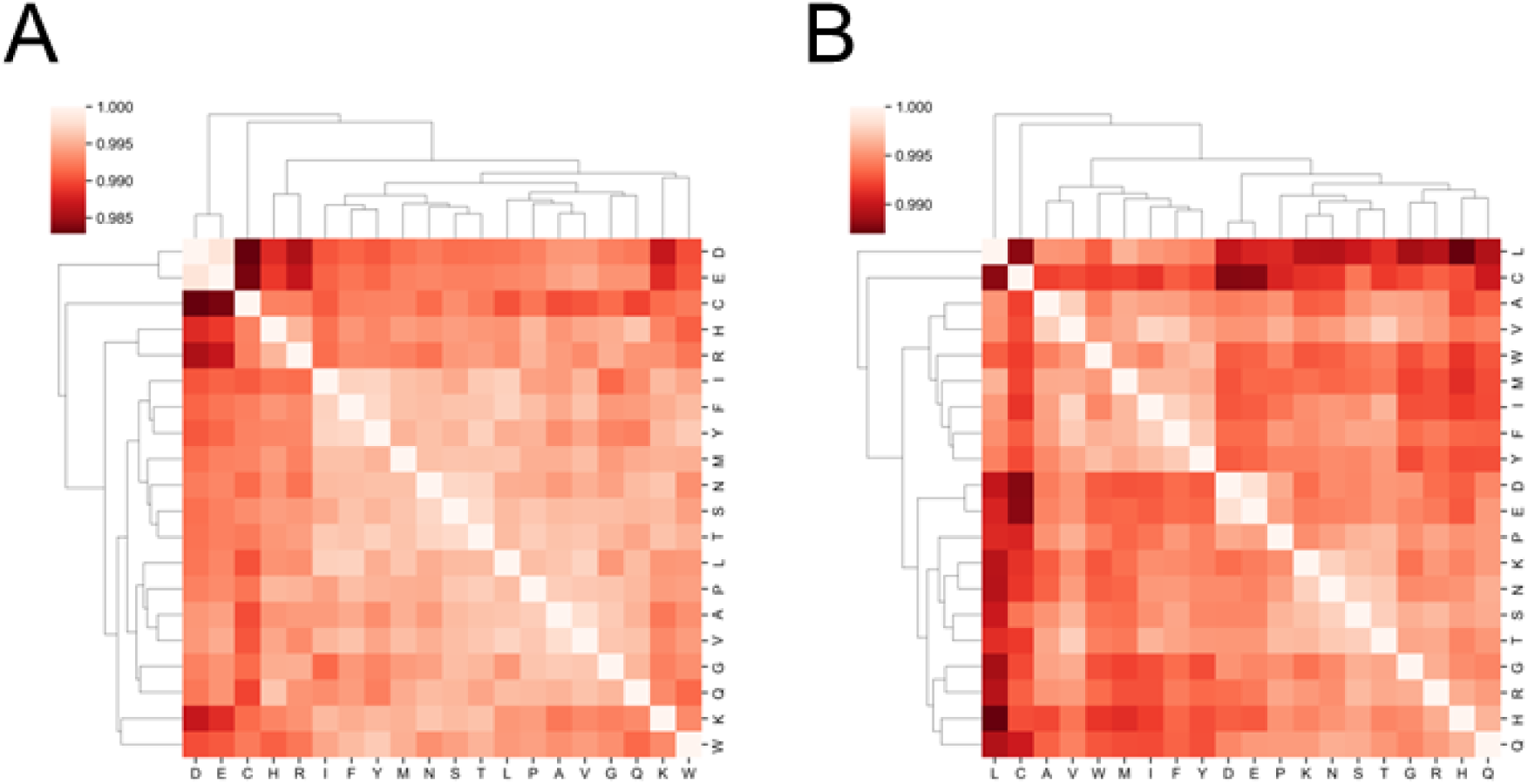
Hierarchical clustering of vector similarities for single-point mutations. **(A, B)** Pairwise cosine similarity matrices for peptide vectors derived from all 20 amino acid substitutions at position 2 (**A**) and position 6 (**B**) of the reference sequence. The heatmap color scale represents the degree of similarity between mutant vectors, ranging from identity (white, 1.000) to lower similarity (dark red). Dendrograms along the axes depict the hierarchical clustering of amino acid residues based on the calculated cosine distances between their corresponding cyclic peptide vectors.

## References

[1] Xiao W, Jiang W, Chen Z, Huang Y, Mao J, Zheng W, Hu Y, Shi J (2025) Advance in peptide-based drug development: delivery platforms, therapeutics and vaccines. Sig Transduct Target Ther 10:. 10.1038/s41392-024-02107-5

[2] Rossino G, Marchese E, Galli G, Verde F, Finizio M, Serra M, Linciano P, Collina S (2023) Peptides as Therapeutic Agents: Challenges and Opportunities in the Green Transition Era. Molecules 28:7165. 10.3390/molecules28207165

[3] Ji X, Nielsen AL, Heinis C (2024) Cyclic Peptides for Drug Development. Angew Chem Int Ed 63:. 10.1002/anie.202308251

[4] Lamers C (2022) Overcoming the Shortcomings of Peptide-Based Therapeutics. Future Drug. Discov. 4:. 10.4155/fdd-2022-0005

[5] de la Torre BG, Albericio F (2020) Peptide Therapeutics 2.0. Molecules 25:2293. 10.3390/molecules25102293

[6] Joo S (2012) Cyclic Peptides as Therapeutic Agents and Biochemical Tools. Biomolecules and Therapeutics 20:19–26. 10.4062/biomolther.2012.20.1.019

[7] Lin H, Zhu C, Shang T, Zhu N, Lin K, Zhang C, Shao X, Wang X, Duan H (2025) HighPlay: Cyclic Peptide Sequence Design Based on Reinforcement Learning and Protein Structure Prediction. J. Med. Chem. 68:12047–12057. 10.1021/acs.jmedchem.5c00896

[8] Li Q, Vlachos EN, Bryant P (2025) Design of linear and cyclic peptide binders from protein sequence information. Commun Chem 8:. 10.1038/s42004-025-01601-3

[9] Rettie SA, Juergens D, Adebomi V, Bueso YF, Zhao Q, Leveille AN, Liu A, Bera AK, Wilms JA, Üffing A, Kang A, Brackenbrough E, Lamb M, Gerben SR, Murray A, Levine PM, Schneider M, Vasireddy V, Ovchinnikov S, Weiergräber OH, Willbold D, Kritzer JA, Mougous JD, Baker D, DiMaio F, Bhardwaj G (2025) Accurate de novo design of high-affinity protein-binding macrocycles using deep learning. Nat Chem Biol 21:1948–1956. 10.1038/s41589-025-01929-w

[10] Lin Z, Akin H, Rao R, Hie B, Zhu Z, Lu W, Smetanin N, Verkuil R, Kabeli O, Shmueli Y, Dos Santos Costa A, Fazel-Zarandi M, Sercu T, Candido S, Rives A (2023), Evolutionary-scale prediction of atomic-level protein structure with a language model. Science 379, 1123–1130. 10.1126/science.ade2574

[11] Frank M, Ni P, Jensen M, Gerstein MB (2024) Leveraging a large language model to predict protein phase transition: A physical, multiscale, and interpretable approach. Proc. Natl. Acad. Sci. U.S.A. 121:. 10.1073/pnas.2320510121

[12] McInnes L, Healy J, Melville J (2020) UMAP: Uniform Manifold Approximation and Projection for Dimension Reduction. arXiv:1802.03426v3. 10.48550/arXiv.1802.03426

[13] Mariani V, Biasini M, Barbato A, Schwede T (2013) lDDT: a local superposition-free score for comparing protein structures and models using distance difference tests. Bioinformatics 29:2722–2728. 10.1093/bioinformatics/btt473

[14] Zhang Y (2005) TM-align: a protein structure alignment algorithm based on the TM-score. Nucleic Acids Research 33:2302–2309. 10.1093/nar/gki524

[15] Rettie SA, Campbell KV, Bera AK, Kang A, Kozlov S, Bueso YF, De La Cruz J, Ahlrichs M, Cheng S, Gerben SR, Lamb M, Murray A, Adebomi V, Zhou G, DiMaio F, Ovchinnikov S, Bhardwaj G (2025) Cyclic peptide structure prediction and design using AlphaFold2. Nat Commun 16:. 10.1038/s41467-025-59940-7

[16] Wu Y, Zhang N, Hashimoto K, Xia C, Dijkstra JM (2021) Structural Comparison Between MHC Classes I and II; in Evolution, a Class-II-Like Molecule Probably Came First. Front. Immunol. 12:. 10.3389/fimmu.2021.621153.

[17] Lipinski CA, Lombardo F, Dominy BW, Feeney PJ (1997) Experimental and computational approaches to estimate solubility and permeability in drug discovery and development settings. Advanced Drug Delivery Reviews 23:3–25. 10.1016/S0169-409X(96)00423-1

[18] Dobson CM (2004) Chemical space and biology. Nature 432:824–828. 10.1038/nature03192

